# Ciliary biology intersects autism and congenital heart disease

**DOI:** 10.1101/2024.07.30.602578

**Authors:** Nia Teerikorpi, Micaela C. Lasser, Sheng Wang, Elina Kostyanovskaya, Ethel Bader, Nawei Sun, Jeanselle Dea, Tomasz J. Nowakowski, A. Jeremy Willsey, Helen Rankin Willsey

## Abstract

Autism spectrum disorder (ASD) commonly co-occurs with congenital heart disease (CHD), but the molecular mechanisms underlying this comorbidity remain unknown. Given that children with CHD come to clinical attention by the newborn period, understanding which CHD variants carry ASD risk could provide an opportunity to identify and treat individuals at high risk for developing ASD far before the typical age of diagnosis. Therefore, it is critical to delineate the subset of CHD genes most likely to increase the risk of ASD. However, to date there is relatively limited overlap between high confidence ASD and CHD genes, suggesting that alternative strategies for prioritizing CHD genes are necessary. Recent studies have shown that ASD gene perturbations commonly dysregulate neural progenitor cell (NPC) biology. Thus, we hypothesized that CHD genes that disrupt neurogenesis are more likely to carry risk for ASD. Hence, we performed an *in vitro* pooled CRISPR interference (CRISPRi) screen to identify CHD genes that disrupt NPC biology similarly to ASD genes. Overall, we identified 45 CHD genes that strongly impact proliferation and/or survival of NPCs. Moreover, we observed that a cluster of physically interacting ASD and CHD genes are enriched for ciliary biology. Studying seven of these genes with evidence of shared risk (*CEP290, CHD4, KMT2E, NSD1, OFD1, RFX3, TAOK1*), we observe that perturbation significantly impacts primary cilia formation *in vitro*. While *in vivo* investigation of *TAOK1* reveals a previously unappreciated role for the gene in motile cilia formation and heart development, supporting its prediction as a CHD risk gene. Together, our findings highlight a set of CHD risk genes that may carry risk for ASD and underscore the role of cilia in shared ASD and CHD biology.

## INTRODUCTION

Autism spectrum disorders (ASD) are complex neurodevelopmental conditions that commonly co-occur with congenital heart disease (CHD) (Bean Jaworski et al. 2017; Marino et al. 2012). For example, a CHD diagnosis increases the likelihood of an ASD diagnosis approximately 2-fold (Gu et al. 2023). Both ASD and CHD are highly heritable and share genetic risk (Homsy et al. 2015; Jin et al. 2017; S. Zaidi et al. 2013; A. J. Willsey et al. 2018; De Rubeis et al. 2014; Satterstrom et al. 2020) and genes impacted by rare likely gene disrupting variants in both conditions are 15-fold more likely to be annotated as chromatin modifiers (Jin et al. 2017). Our group, using joint network propagation of ASD and CHD genes, previously identified significant overlap of associated molecular networks, and pinpointed chromatin modification, NOTCH signaling, MAPK signaling, and ion transport as potential areas of shared biology (Rosenthal et al. 2021). Taken together, there is strong evidence that ASD and CHD likely share common biology, yet the relevant molecular mechanisms that underlie comorbidity remain unclear.

Because CHD is generally identified by the newborn stage, the co-morbidity between CHD and ASD affords a potentially critical opportunity to ascertain ASD patients, conduct observational studies, and begin therapeutic interventions far sooner than typically possible (A. J. Willsey et al. 2018; Homsy et al. 2015; Jin et al. 2017). This strategy will be most effective if CHD patients can be stratified by risk of developing ASD, yet we do not currently have the ability to do this, due to the relatively limited overlap between high confidence genes identified in rare-variant based whole exome sequencing studies of ASD and of CHD (Satterstrom et al. 2020; Jin et al. 2017). Therefore, we employed a multiplexed *in vitro* genetic perturbation screen to prioritize CHD genes likely to increase the risk of ASD.

*In vivo* and *in vitro* studies have repeatedly identified neural progenitor cell (NPC) proliferation as a convergent phenotype in ASD (Sun et al. 2024; H. R. Willsey et al. 2022, 2021; Sacco, Cacci, and Novarino 2018; Packer 2016; Iakoucheva, Muotri, and Sebat 2019; Courchesne et al. 2019; Marchetto et al. 2017; Lalli et al. 2020). We hypothesized that the subset of CHD genes that disrupt neurogenesis will likely carry risk for ASD. Therefore, we performed a multiplexed CRISPR interference (CRISPRi) proliferation and survival screen (Tian et al. 2019) in human NPCs, targeting ASD and CHD genes. Overall, we identified 45 CHD genes that strongly impact NPC biology, as well as, a cluster of ASD and CHD genes that impact NPC proliferation and are putatively involved in ciliary biology. Within this cluster, we showed that all seven genes predicted to share risk for both ASD and CHD (*CEP290, CHD4, KMT2E, NSD1, OFD1, RFX3, TAOK1*) are required for primary cilia biology in human cells *in vitro*. We also demonstrated that loss of *TAOK1* impairs motile cilia, as well as heart and brain development *in vivo* in *Xenopus*. Together, these results outline a set of CHD genes likely to carry risk for ASD and suggest a role for cilia at the intersection of ASD and CHD shared biology. A finding consistent with recent results implicating tubulin biology in ASD (H. R. Willsey et al. 2022; Sun et al. 2024).

## RESULTS

### Pooled proliferation/survival screen of ASD and CHD genes in NPCs

Previous research has shown that ASD risk gene variants commonly perturb NPC biology (Sun et al. 2024; H. R. Willsey et al. 2022, 2021; Sacco, Cacci, and Novarino 2018; Packer 2016; Iakoucheva, Muotri, and Sebat 2019; Courchesne et al. 2019; Marchetto et al. 2017). Therefore we leveraged a bulk CRISPRi screening approach (Tian et al. 2019) to determine whether some CHD genes also impact the survival and/or proliferation of NPCs. We generated a pooled lentiviral sgRNA library targeting 62 high-confidence ASD genes, 195 CHD genes, and 104 ‘ASD-CHD’ shared risk genes, using at least 5 sgRNAs per gene (361 total genes, **Figure 1A**, see **Methods**). We also included 255 distinct non-targeting control sgRNAs in our library. Next, we generated NPCs from the Allen Institute for Cell Science (AICS) dCAS9 iPSC line, which enables stable CRISPRi in iPSC-derived neuronal lines (Sun et al. 2024; Tian et al. 2019). We transduced these NPCs with the pooled lentiviral sgRNA library and then passaged them for 20 days. We collected cells at day 0, 5, 10, and 20, and determined the number of cells expressing each sgRNA at each timepoint by sequencing the sgRNA protospacer, which is the unique sequence that targets the sgRNA to a specific gene or identifies the non-targeting control sgRNAs (**Figure 1B**).

**Figure 1.**
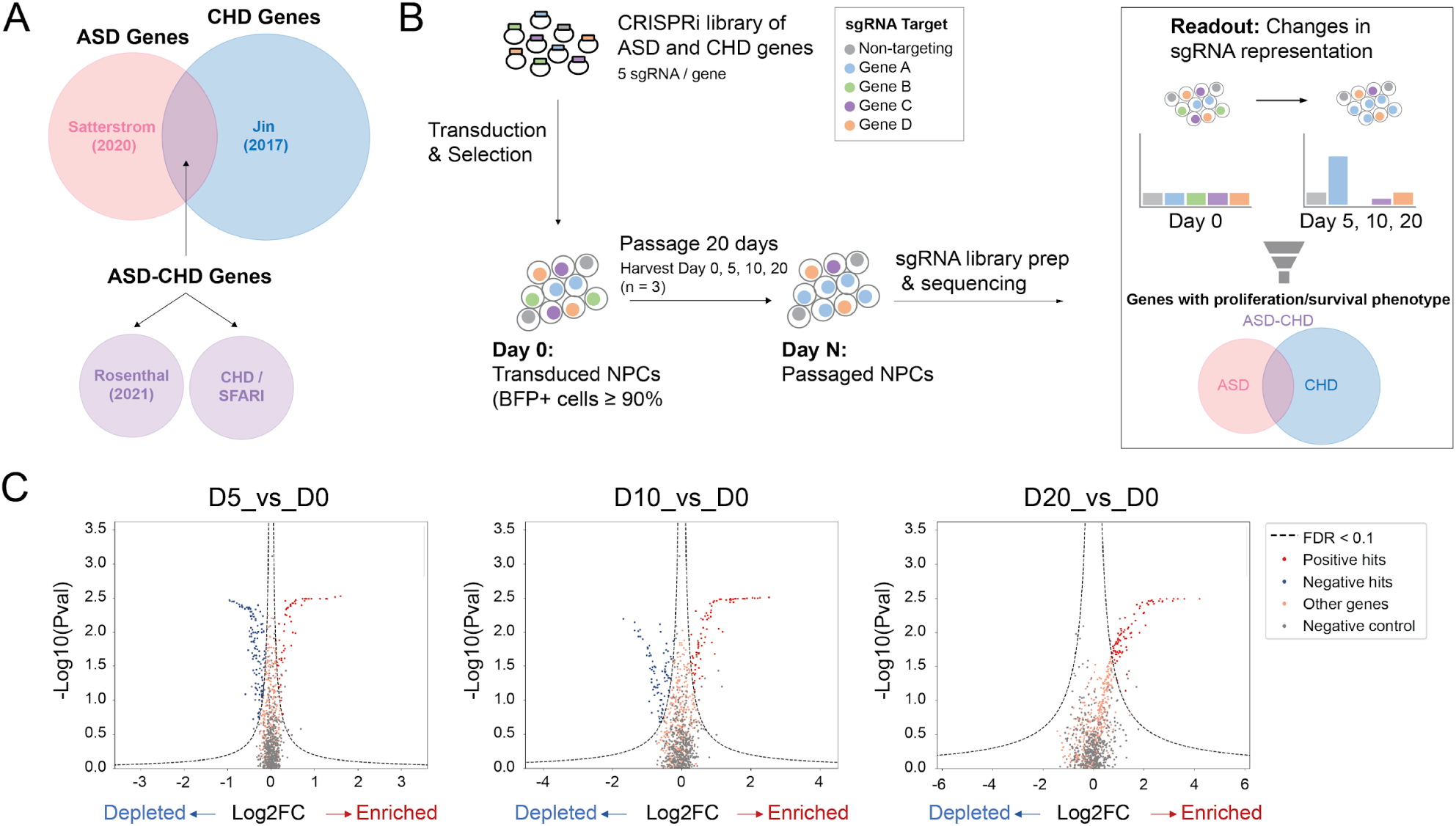
Pooled Proliferation/Survival Screen of ASD and CHD Genes in NPCs. (A) Venn diagram showing the origin of genetic targets. ASD genes (pink; only identified in Satterstrom et al. 2020), CHD genes (blue; only identified in Jin et al. 2017), and ASD-CHD genes (purple). ASD-CHD genes (1) share genetic risk (Satterstrom et al. 2020, Jin et al. 2017), (2) have predicted shared risk (Rosenthal et al. 2021); or (3) are CHD genes (Jin et al. 2017) present in the SFARI Gene database. (B) Strategy for CRISPRi Screen. Cells are harvested at Day 0, 5, 10, and 20 and then sgRNA representation at Day 5, 10, and 20 is compared against Day 0. (C) Volcano plots summarizing knock-down phenotypes and statistical significance (Mann-Whitney U test) for sgRNAs in the pooled screen. Dashed lines: cutoff for hit sgRNAs (FDR = 0.1) represented by both red (significantly enriched) and blue (significantly depleted) circles. Gray circles represent non-targeting RNA and orange circles represent non-significant genes. See also Table S1.

We used the MAGeCK bioinformatics pipeline (W. Li et al. 2014; Tian et al. 2019) to compare sgRNA representation at each timepoint versus day 0 and calculate gene-level fold-changes and false discovery rates (FDRs) as per the methods in Tian *et al*. (2019). Overall, we identified 24 ASD, 77 CHD, and 44 ASD-CHD genes that impact survival and/or proliferation of NPCs positively or negatively when disrupted (FDR < 0.1, 145 total genes, **Figure 1C, Table S1**), supporting our hypothesis that a subset of CHD genes will disrupt neurogenesis.

### Enrichment of ciliary biology among ASD and CHD genes

To subset the 145 significant genes into groups that potentially represent congruent biological processes, we focused on the 54 genes (9 ASD, 28 CHD, 17 ASD-CHD) with a fold-change in guide representation greater than 1.5x at one or more timepoints (log2FC ≥ 0.585, **Table S2**) and performed k-means clustering (**Figure 2A**). While there are two obvious clusters (enriched vs. depleted sgRNAs), we estimated an ideal cluster number of four using the elbow plot method (**Figure S1**). We observed that all clusters contained ASD, CHD, and ASD-CHD genes, suggesting that ASD and CHD genes converge onto *in vitro* “neurogenesis” phenotypes with similar trajectories. We next aimed to identify the subset of clusters with more connections between genes than predicted by chance, which would suggest an underlying shared biology. To do this, we queried StringDB, a database of known and predicted physical and functional interactions (Szklarczyk et al. 2021). This analysis identified Cluster 1 as containing the only gene set with a significant enrichment of interactions contained in the full StringDB network as well as of protein-protein interactions only (**Figure 2B-C, S2**).

**Figure 2.**
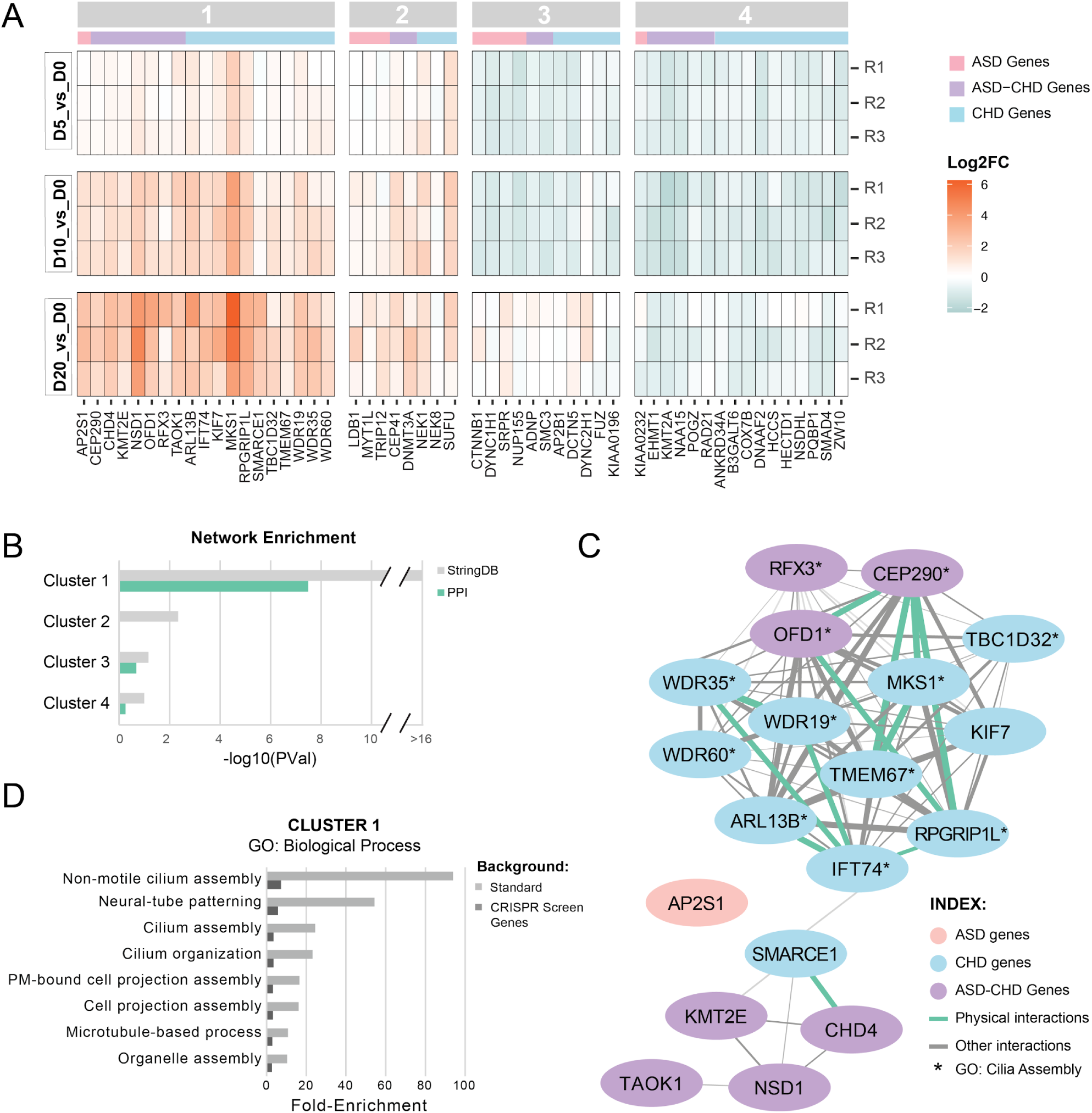
A subset of ASD and CHD genes converge on cilia biology. (A) Heat-map showing gene knock-downs grouped by k-means clustering (see Figure S1). R1-3 represent biological replicates. Cutoff for genes: p-value < 0.05, absolute value of log2(fold-change) ≥ 0.585 for at least one time point. ASD genes are denoted with a pink bar, CHD genes with a blue bar, and ASD-CHD genes with a purple bar., (B) Genes within K-means cluster 1 are more connected than expected by chance based on interactions from StringDB. Enrichment for clusters 1-4 was calculated for the all interactions inStringDB (gray) and for just the physical interactions in StringDB (green). P-values are not corrected for multiple comparisons. (C) Visualization of the network of cluster 1 genes, built from the interactions cataloged in StringDB. ASD genes are in pink, CHD genes are in blue, and ASD-CHD genes are in purple. Interactions are represented by a green line (physical interactions) or gray line (Other StringDB interaction categories). (D) ToppGene GO enrichment analysis (Biological Process) of genes from cluster 1 shows enrichment of ciliary pathways, for both the standard background (light gray) and a custom background consisting of the 361 genes screened here (dark gray). Only terms with a false discovery rate less than 0.05 (Benjamini-Hochberg procedure) on both backgrounds are displayed. See also Figures S1-S2, Tables S2-S3.

To identify potential biological pathways or processes underlying Cluster 1, we utilized ToppGene (J. Chen et al. 2009) to perform gene ontology enrichment analysis of the Cluster 1 genes, using either the standard background set of genes or, more stringently, only the 361 genes screened here as background. We observe significant enrichment of only eight Biological Process terms across both backgrounds (Benjamini-Hochberg FDR < 0.05, **Figure 2D**), with three of the four highest fold-enrichments related to cilia, microtubule-based hair-like organelles critical for proliferation, patterning, signaling, and survival (D. Zaidi, Chinnappa, and Francis 2022; Mill, Christensen, and Pedersen 2023; Anvarian et al. 2019). “Neural-tube patterning”, a process reliant on cilia, rounds out the top four most strongly enriched terms (Guemez-Gamboa, Coufal, and Gleeson 2014). We similarly observed significant enrichment of eight Cellular Component terms, with all eight terms relating to cilia and microtubule biology (**Table S3**). These results suggest that genes in Cluster 1, when perturbed, could impact cilia, leading to changes in both brain and heart development (Guemez-Gamboa, Coufal, and Gleeson 2014; Djenoune et al. 2022; Shaikh Qureshi and Hentges 2024; Youn and Han 2018). Finally, while overrepresented sgRNAs as a group are enriched for cilia-related genes, this enrichment is predominantly driven by Cluster 1 genes (Cluster 2 genes alone are not significantly enriched for any cilia-related gene sets and the fold-enrichments for significant GO terms are lower when combining Cluster 1 and 2).

### ASD-CHD shared genes are required for ciliary biology

Based on the enrichment of Cluster 1 genes for GO terms related to ciliary biology, we next directly tested whether individual disruption of a subset of these genes results in cilia defects. More specifically, we selected the seven ASD-CHD shared risk genes (*CEP290, CHD4, KMT2E, NSD1, OFD1, RFX3, TAOK1*), as these genes are most likely to represent shared biology. Of these genes, four (*CEP290*, *CHD4*, *OFD1*, and *RFX3*) have previously described roles in ciliary biology (Rachel et al. 2015; Wu et al. 2020; Morleo, Pezzella, and Franco 2023; B. Chen et al. 2018; Robson et al. 2019; Marley and von Zastrow 2012). In contrast, the remaining three (*KMT2E*, *NSD1*, and *TAOK1*) have not been directly implicated in ciliary biology. That being said, *KMT2E* and *TAOK1* are known to regulate microtubules (Zhao et al. 2016; Draviam et al. 2007), the main cytoskeletal component of cilia.

To determine the extent to which these ASD-CHD shared risk genes play a role in ciliary biology, we individually disrupted expression of the ASD-CHD genes listed above using CRISPRi in mitotically-arrested retinal pigment epithelial cells (RPE1), a robust *in vitro* model for evaluating primary cilia dynamics (May-Simera et al. 2018). All seven genes are expressed in these cells and we confirmed strong CRISPRi knock-downs by qPCR (**Figure S3A**). We first examined the percentage of ciliated cells and observed that repression of each of these seven ASD-CHD genes resulted in significant decreases in the percentage of ciliated cells when compared to two independent, non-targeting control knock-downs (**Figure 3A-B, S3B, S4A)**. As an orthogonal assay of cilia disruption, we assessed whether individual repression of these genes impacted cilia length in mitotically-arrested cells. Length measurements provide insight into both stability and functionality of cilia, which rely on length changes for various cellular processes, such as cell cycle regulation (Avasthi and Marshall 2012; D. Zaidi, Chinnappa, and Francis 2022). With the exception of *CHD4*, repression of each of the genes resulted in decreased cilia length (**Figure 3C-D, S3C, S4B)**. Taken together, these data provide strong evidence that all seven of these shared ASD-CHD genes intersect ciliary biology.

**Figure 3.**
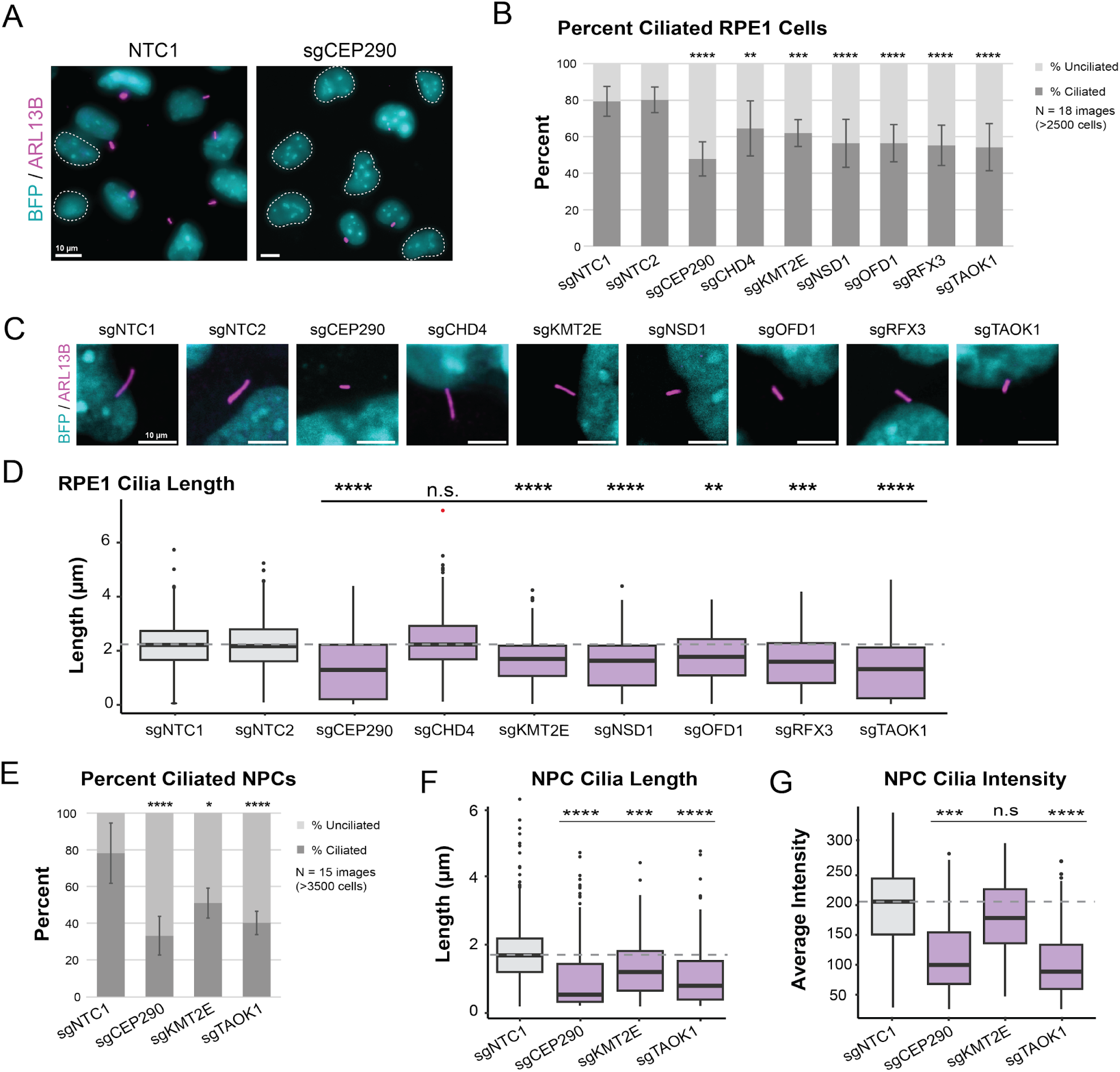
Knock-down of ASD-CHD genes disrupts primary cilia. (A) Representative image of ciliated RPE1 cells transduced (BFP+) with a non-targeting control sgRNA (sgNTC1) and a *CEP290* targeting sgRNA (sgCEP290). BFP+ cells (cyan) without primary cilia are circled with a white dotted line. Cilia are labeled by ARL13B (magenta). (B) Repression of ASD-CHD genes (*CEP290, CHD4, KMT2E, NSD1, OFD1, RFX3, TAOK1*) results in a decrease in percent ciliated cells. We quantified percent ciliated cells in ≥2500 RPE1 cells across 18 images (3 biological replicates) using CiliaQ(ref). (C) Representative image of cilia length in RPE1 cells for ASD-CHD knock-downs. BFP+ cells (cyan) and ARL13B (magenta). (D) We observed a decrease in primary cilia length for select ASD-CHD gene knock-downs. We captured ≥250 cells across 15 images (3 biological replicates) for quantification of cilia length (µm) in RPE1 cells using CiliaQ (Hansen et al. 2021). (E) Percent cilia phenotypes are replicated in NPCs for a subset of ASD-CHD gene knock-downs (*CEP290, KMT2E, TAOK1*). We captured ≥3500 cells across 15 images (3 biological replicates) for quantification of percent ciliated cells in NPCs using CiliaQ (Hansen et al. 2021). (F) Cilia length phenotypes were also replicated in NPCs. We captured ≥350 cells across 15 images (3 biological replicates) and measured length using CiliaQ (Hansen et al. 2021). (G) In addition to decreased cilia length, we also observed a decrease in ARL13B signal for *CEP290* and *TAOK1* knock-down NPCs. Using the images from (F), we measured ARL13B intensity (a.u.) for quantification using CiliaQ (Hansen et al. 2021). All data were normalized based on average cell density of the non-targeting control sgRNA. *Significance (Dunn’s multiple comparisons)*: *p < 0.05; \*\**p* < 0.01; ***p < 0.001; \*\*\*\**p* < 0.0001; n.s., not significant (*p* > 0.05). See also Figures S3-S4.

Next, to more directly assess whether cilia function during brain development may be compromised, we selected three of these genes (*CEP290*, *KMT2E*, and *TAOK1*) for experiments in human iPSC-derived NPCs. These genes represent different degrees of evidence for ciliary relevance and ASD/CHD association. *CEP290* is a well-characterized CHD gene (Jin et al. 2017) present in the SFARI list of ASD genes (Category 2S) with a known role in ciliary biology as a component of the ciliary basal body, the modified centrosome that organizes cilia formation (Rachel et al. 2015; Firat-Karalar 2018; Wu et al. 2020). In contrast, *TAOK1* and *KMT2E* are ASD genes (Satterstrom et al. 2020; Fu et al. 2022) with predicted risk for CHD based on network propagation (Rosenthal et al. 2021), but have no known role at the cilium. As in RPE1 cells, we assessed the percentage of ciliated cells as well as cilia length and observed significant alterations of both after knock-down of each of these three genes (**Figure 3E-F, S4C-D**). Within intact cilia, we also observed a decrease in the intensity of the cilia marker ARL13B for *CEP290* and *TAOK1* knock-down cells (**Figure 3G, S4E**), suggesting that these genes may impact ARL13B expression and/or localization at the cilium. Together, these results demonstrate that these ASD-CHD shared risk genes are required for primary cilia biology in both NPCs and RPE1 cells.

### *TAOK1* depletion disrupts brain and heart development *in vivo*

While *TAOK1* is a high-confidence ASD gene (Satterstrom et al. 2020; Fu et al. 2022), its association with CHD has only been predicted by network propagation (Rosenthal et al. 2021). Therefore, we sought to elaborate its role *in vivo* in heart development and on motile cilia, which have repeatedly been implicated in CHD pathobiology (N. T. Klena, Gibbs, and Lo 2017; Gabriel, Young, and Lo 2021).

First, we expressed GFP-tagged human TAOK1 in *Xenopus laevis* motile multiciliated cells and observed localization at ciliary basal bodies and axonemes, consistent with a potential function at motile cilia (**Figure S5A**). Then we depleted *taok1* in the multiciliated epidermis of diploid *Xenopus tropicalis* embryos using a translation-blocking morpholino (**Figure 4A**) and classified multiciliated cells based on the extent of cilia loss. In Taok1-depleted embryos, we observed that 98% of ciliated cells had a moderate to severe phenotype, which is significantly greater than the 19% observed in the control embryos injected with a non-targeting morpholino (**Figure 4B-C**). We observed the strongest effect size when comparing the rate of cells with a severe phenotype (severe versus no phenotype odds ratio = 338.7, p < 0.0001 by two-sided Fisher’s exact test, moderate versus no phenotype odds ratio = 135.0, p < 0.0001). Our results thus demonstrate that TAOK1 is not only critical for primary cilia *in vitro*, but also for motile cilia *in vivo*.

**Figure 4.**
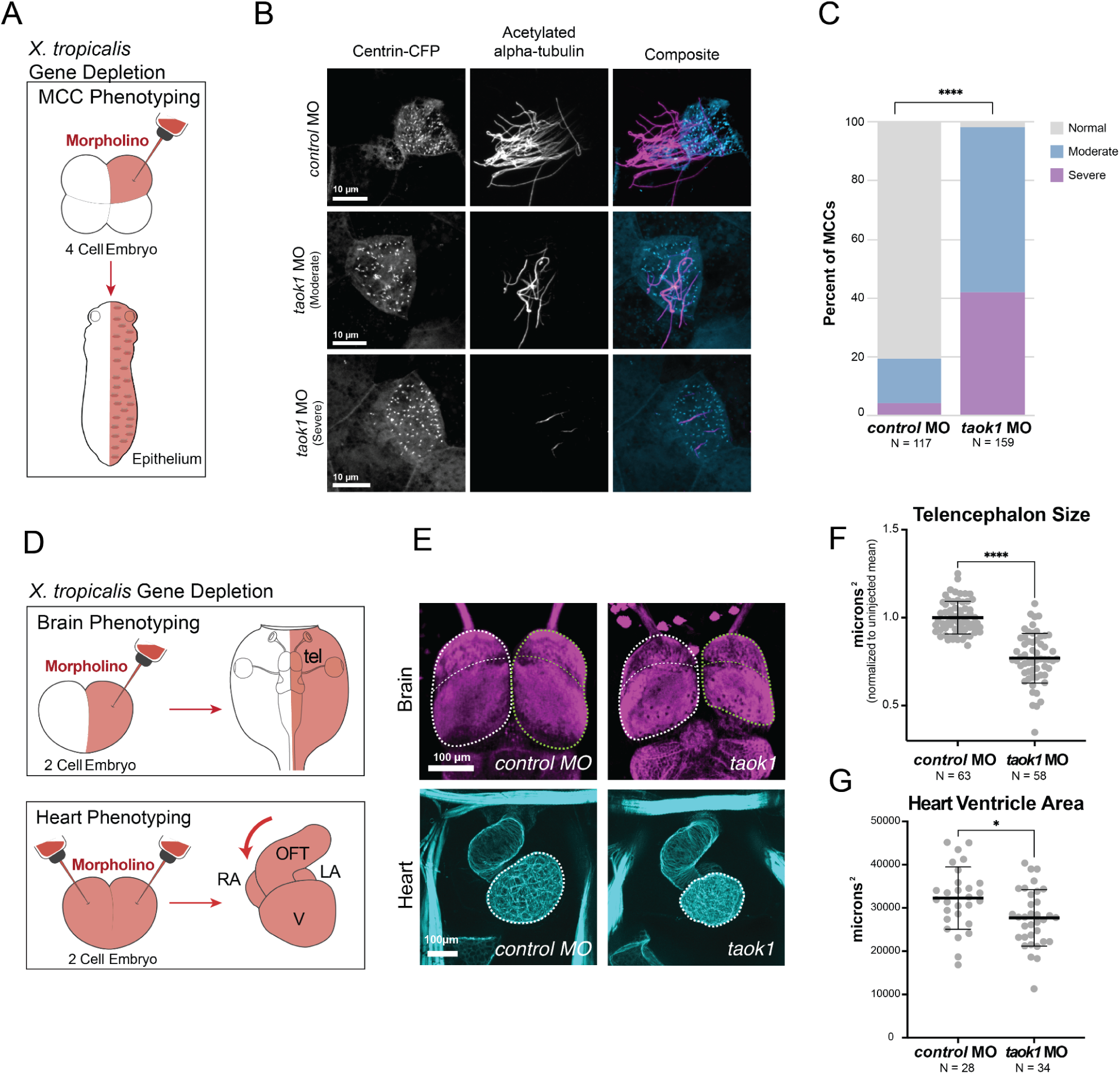
TAOK1 is required for brain and heart development *in vivo*. (A) Depletion strategy. Morpholino and a tracer (Centrin-CFP) are injected at the four-cell stage for phenotyping of epithelial multiciliated cells (MCCs) in *Xenopus tropicalis*. (B) Stage 26 *X. tropicalis* MCCs expressing Centrin-CFP are stained for acetylated α-Tubulin (cilia, magenta). Injection of *taok1* morpholino causes moderate to severe loss in cilia relative to injected non-targeting control. (C) Quantification of cilia phenotype in MCCs by condition. Y-axis shows percent of cilia classified as normal (≥50 cilia, grey), moderate (10-50 cilia, blue), or severe (<10 cilia, purple). *N = X cells across 3 embryos. Significance was calculated using Chi-squared test (****p < 0.00001) (D) Depletion strategy (Rosenthal et al. 2022). Morpholino and a tracer (dextran) are injected at the two-cell stage, either into one cell (brain phenotyping) or both (heart phenotyping). Stage 46 *X. tropicalis* tadpoles are phenotyped for brain (top) anatomy by comparing bilateral symmetry. The tadpoles are phenotyped for heart (bottom) by comparing heart looping direction and ventricle size. Annotations: telencephalon (tel), outflow tract (oft), ventricle (V), right atrium (RA), and left atrium (LA). (E) Images of brain/telencephalon (top, β-tubulin, magenta) and heart (bottom, phalloidin/actin, cyan). Negative control standard morpholino is on the left and *taok1* morpholino is on the right. Brain: Disruption of *taok1* results in decreased telencephalon size (green dotted outline) relative to control. Heart: Loss of *taok1* results in decreased ventricle size relative to the control. Scale bars: 100 µm. (F) Telencephalon size variance for *taok1* depletion. Variation in forebrain size relative to the uninjected side (µm^2) for control morpholino-injected animals versus *taok1* depleted animals. Significance was calculated using a Mann-Whitney rank sum test (\*\*\*\**p* < 0.0001). (G) Ventricle size variance for *taok1* depletion. Variation in ventricle size for *taok1-*depleted embryos versus control injected embryos. Significance was calculated using non-parametric Mann-Whitney rank sum test (\**p* < 0.05). See also Figure S5.

Next, we tested whether disruption of *taok1* leads to brain and heart developmental phenotypes *in vivo*. For the brain, we created unilaterally *taok1*-depleted tadpoles by morpholino injection into one cell of two-cell stage *X. tropicalis* embryos (**Figure 4D**) and phenotyped tadpoles for variation in forebrain size relative to the uninjected side, as previously described for many ASD genes (Rosenthal et al. 2021; H. R. Willsey et al. 2021, 2020, 2018). *Taok1* depletion significantly decreased telencephalon size compared to the control standard morpholino injection (**Figure 4E-F**), a previously-characterized convergent phenotype of ASD gene disruption (H. R. Willsey et al. 2021; Villa et al. 2022; Ye et al. 2015; Sacco, Gabriele, and Persico 2015). For heart phenotyping, we created bilaterally depleted embryos by morpholino injection into both blastomeres at the two-cell stage (**Figure 4D**), and assessed tadpole hearts for structural abnormalities as previously described for CHD risk genes in *Xenopus* (Garfinkel and Khokha 2017; Duncan and Khokha 2016; Rosenthal et al. 2021). We observed a decrease in heart ventricle size in *taok1*-depleted tadpoles relative to animals injected with a control standard morpholino (**Figure 4E-G**). These results suggest that *TAOK1* is required for heart development, reinforcing the putative role in ciliary biology hypothesized here, as cilia defects underlie many CHDs (Yuan, Zaidi, and Brueckner 2013; Fakhro et al. 2011; Gabriel, Young, and Lo 2021; Jin et al. 2017; N. T. Klena, Gibbs, and Lo 2017; Y. Li et al. 2015; Djenoune et al. 2022; N. Klena et al. 2016). Additionally, we identified several patients with *TAOK1 de novo* variants in the DECIPHER database that present with both ASD and structural congenital heart abnormalities (Hunter et al. 2022), consistent with the possibility that *TAOK1* is in fact a shared risk gene, as predicted by (Rosenthal et al. 2021). Together, our work demonstrates that *TAOK1* is required for primary and motile cilia, heart development, and brain development in *Xenopus*, and supports its classification as a bona fide risk gene for both CHD and ASD.

## DISCUSSION

In conclusion, we identified ASD, CHD, and predicted shared risk genes that impact human neural progenitor cell proliferation and/or survival, thereby prioritizing a subset of CHD genes that may carry risk for ASD. Furthermore, we presented both *in vitro* and *in vivo* evidence implicating cilia biology at the intersection of ASD and CHD genetics. More specifically, we identified 24 ASD, 77 CHD, and 44 ASD-CHD genes that perturb NPC biology. In a subset of these genes, we observed significant enrichment of GO terms related to ciliary biology. We validated this finding, showing that repression of seven ASD-CHD predicted shared risk genes individually (*CEP290, CHD4, KMT2E, NSD1, OFD1, RFX3, TAOK1*) all led to primary cilia defects *in vitro* in human cells. Finally, we identified an additional role for *TAOK1* in motile cilia as well as heart and brain development *in vivo* in *Xenopus*, supporting the prediction that it carries risk for CHD. These findings are consistent with recent work identifying TAOK1 as a predicted regulator of TTBK2, a known ciliary regulator (Loukil, Barrington, and Goetz 2021; Bashore et al. 2023)..

Building on prior studies identifying enrichment in chromatin regulation, NOTCH signaling, and MAPK signaling for genes with shared risk for ASD and CHD (Rosenthal et al. 2021; D. Zaidi, Chinnappa, and Francis 2022; S. Zaidi et al. 2013; Fakhro et al. 2011), our study adds ciliary biology as a point of vulnerability intersecting these disorders. Ciliary underpinnings in CHD have been well-established (Yuan, Zaidi, and Brueckner 2013; Fakhro et al. 2011; Gabriel, Young, and Lo 2021; Jin et al. 2017; N. T. Klena, Gibbs, and Lo 2017; Y. Li et al. 2015; Djenoune et al. 2022; N. Klena et al. 2016), but their implications for our understanding of ASD are less commonly appreciated despite some evidence in the literature (Marley and von Zastrow 2012; H. R. Willsey et al. 2020, 2018; Frasca et al. 2020; Rosengren et al. 2018; Di Nardo et al. 2020). This work, combined with our group’s recent work showing that ASD-associated chromatin regulators also regulate microtubules (Lasser et al. 2023), sheds new light on the previous identification of shared ASD-CHD biology around chromatin regulation. It is possible that the finding of shared ontology around chromatin regulation may actually represent signals related to microtubule biology, and, therefore, ciliary biology since microtubules are the major structural component of cilia. Other recent work from our group also shows broader convergence of ASD genes onto microtubule biology (Sun et al. 2024), suggesting this ciliary finding may be more broadly applicable to ASD mechanisms beyond just comorbidity with CHD.

Cilia have diverse cellular functions, including regulating cell proliferation, differentiation, and even neuronal excitability (Mill, Christensen, and Pedersen 2023; Malicki and Johnson 2017; Mitchison and Valente 2017; Tereshko et al. 2021). Conversely, defects in cell proliferation can affect cilia formation (Kasahara and Inagaki 2021), obscuring which process was primarily affected in our NPC screen. However, we also showed that these knock-downs caused alterations in cilia length in mitotically-arrested cells, supporting a direct role at cilia. Nevertheless, general defects in microtubule stability will affect both cell proliferation/survival via the mitotic spindle and cilia formation and length, so it’s unclear which of these processes are central to the phenotypes observed here (and in patients). These effects are also commonly dose-dependent. Therefore future work could be focused to untangle these processes, for example in post-mitotic cells, and with respect to gene dosage, using weaker gene repression than we used here. Cilia are also the sole cellular site of sonic hedgehog (SHH) signaling (Bangs and Anderson 2017), so the effect of ASD/CHD gene perturbation on the associated process of differentiation and the associated signaling cascades during heart and brain development is an exciting future direction (H. R. Willsey et al. 2021). Finally, cilia have cell type-specific functions, so it will be important for future work to explore how these genes affect cilia formation and function in a cell type-dependent manner.

Overall, our research generates insights into the shared biology underlying ASD and CHD, suggests a class of genes that are likely to carry risk for both conditions, and provides a path forward for investigating known and predicted risk genes during heart and brain development.

## MATERIALS AND METHODS

### Human cell culture

#### Human iPSCs

Allen Institute for Cell Science (AICS) BFP-tagged dCas9-KRAB WTC iPSC line (AICS-0090-391, MONO-ALLELIC TagBFP-TAGGED dCas9-KRAB WTC) were cultured in mTESR Plus Medium (Stem Cell Technologies; Cat. No. 05825) on Matrigel (Fisher Scientific; Cat. No. 08-774-552) coated Cell Culture Dishes (Corning; Cat. No. 08-774-552) diluted in DMEM F12 (Fisher Scientific; Cat. No. 11320-082). mTESR Plus Medium was replaced every day and cells (70-90% confluent) were passaged using Accutase (Stem Cell Technologies; Cat. No. 07920), then re-plated in mTESR Plus Medium with the addition of 10nM Y-27632 dihydrochloride ROCK inhibitor (Tocris; Cat. No. 125410) for 24 hr.

#### Human iPSC-Derived Neural Progenitor Cells (NPCs)

We generated neural progenitor cells (NPCs) from the AICs dCAS9 iPSC line using a modified version of a monolayer dual-SMAD inhibition protocol combined with small molecules, producing >98% PAX6+ cells (Sun et al. 2024). Briefly, we treated cells with LDN193189, SB431542, and XAV939 for 6 days. The cells were then passaged and cultured with XAV939 alone for 2 more days to generate NPCs. NPCs were then maintained in N2/B27 medium (DMEM F-12, 1x B27 -Vit.A, 1x N2, 1x GlutaMAX, 1x MEM-NEAA, 10ng/ml EGF, 10ng/ml FGF2). The medium was changed every other day and cells were passaged at ∼90% confluence using Accutase.

#### Human RPE1/LentiX-293T

Immortalized hTERT dCas9 RPE1 (Jost et al. 2017) cells were cultured in DMEM F12 (Thermo Fisher Scientific; Cat. No. 11320-082) supplemented with 10% FBS on Corning cell culture dishes. LentiX-293T cells were cultured in DMEM (Fisher Scientific; Cat. no. 10-566-024) supplemented with 1x MEM-NEAA, and 10% FBS. Both RPE1s and 293Ts were passaged using 0.25% trypsin-EDTA.

### Pooled Proliferation/Survival Screen

Guides were designed using the CRISPRiaDesign tool (https://github.com/mhorlbeck/CRISPRiaDesign; (Horlbeck et al. 2016). We selected 100 high-confidence ASD-risk (Satterstrom et al. 2020) and 248 CHD-risk (Jin et al. 2017) genes from studies that have leveraged the statistical power of recurrent rare de novo variants in ASD probands. We also identified 104 shared risk ‘ASD-CHD’ genes defined by having at least one of the following characteristics: (1) are present in both ASD and CHD gene lists (Satterstrom et al. 2020; Jin et al. 2017), (2) are CHD genes (Jin et al. 2017) found in the SFARI Gene Database (Gene score: 1-2, Syndromic), or (3) were predicted to share risk by network proximity analysis (Rosenthal et al. 2021). Due to overlap between these three gene sets (ASD, CHD, ASD-CHD), we ultimately designed a library targeting 361 total genes (62 high-confidence ASD genes, 195 CHD genes, and 104 ‘ASD-CHD’ shared risk genes). We designed five sgRNAs per gene and selected 255 non-targeting control sgRNAs (10% total sgRNA). Guides were cloned into pMK1334 (CROPseq-Guide-Puro vector (Tian et al. 2019), RRID:Addgene_ 127965; gifted by Martin Kampmann). We then screened for correctly assembled clones by colony PCR and further validated them using Sanger sequencing. The library balance of sgRNA sequences were then assessed and verified by Ion Torrent Sequencing.

To produce lentivirus for our CRISPRi library, we used LentiX-293T (Clontech) that were maintained in DMEM with Glutamax (Fisher Scientific, Cat. No. 10566016), MEM-NEAA (Fisher Scientific; Cat. no. 11140-050) and 10% FBS. Lentiviral packaging was performed by seeding 2.4 million cells per 10 cm dish, then transfecting with 2.5 µg equimolar packaging mix (pMDL, pRSV, pVSV-g), 2.5 µg sgRNa vector (PMK1334) using OptiMEM and Lipofectamine 2000 (Fisher Scientific; Cat. no. 11668019). 72 hours later, we collected the supernatant, filtered with a 0.45 µm PVDF syringe filter, and concentrated the virus using the Lenti-X Concentrator (Takara Bio, Cat. No. 631231). HEK293 media was replaced with DMEM-F12 when concentrating the virus.

The concentrated virus containing the validated sgRNA library was transduced into NPCs through pooled packaging at 10-20% efficiency to ensure one integration event per cell. We seeded 3 million NPCs each onto two matrigel-coated 10 cm dishes and added 100 uL of concentrated virus that had been resuspended in 1 mL of DMEM F12. Two days later, cells were passaged. 1 million cells were taken for FACS sorting on BFP to ensure no more than 20% transduction efficiency, while the rest were re-plated onto three matrigel-coated 10 cm dishes (3 replicates / 3 million cells per plate) in N2/B27 media containing 3 µg/mL puromycin (Fisher Scientific; Cat. no. 501532829) to select for the pMK1334 sgRNA vector. We refreshed selection media daily, then passaged cells on day 3. Approximately 1 million cells were FACs sorted for expression of BFP (≥85%) as a readout of transduction efficiency. Then 5 million cells from each replicate were harvested (Day 0; library representation ∼1,000 cells per sgRNA). The remainder of the cells were re-plated onto three 10 cm cell culture dishes for later time points. We seeded 2 million NPCs per plate and cultured in N2/B27 medium as described previously. The cells were passaged every 3-5 days and approximately 5 million cells from each replicate were then harvested at days 5, 10 and 20.

We isolated genomic DNA from all samples using the Zymo Quick DNA mini-prep Plus Kit (Zymo Research; Cat. no. D4068). The samples were amplified and prepared for sequencing as described previously (Gilbert et al. 2014).

### Pooled Proliferation/Survival Screen - Data Analysis

Data were analyzed using a bioinformatics pipeline, MAGeCK-iNC (MAGeCK including Negative Controls) as previously described (Tian et al. 2019; W. Li et al. 2014). Briefly, to determine sgRNA counts in each sample, we cropped and aligned the raw sequencing reads to the reference using Bowtie (W. Li et al. 2014). Next, we removed outlier data points (sgRNA count coefficient of variation ≥ 1). Count’s files of timepoints to be compared were then input into MAGeCK to generate log2 fold changes (Log2FC) and p-values for each sgRNA, using the ‘mageck tesk - k’ command. We subtracted the median Log2FC of non-targeting sgRNA from gene-targeting sgRNA to assess changes in gene-targeting sgRNA representation at each timepoint. Gene-level knock-down effects were then determined by taking the mean of individual sgRNA scores for the top 3 sgRNAs targeting a specific gene. Screen positive genes were selected based on a gene-level false discovery rate (FDR) less than 0.1. We then further prioritized genes with a gene-level Log2FC greater than 0.585 for at least one time-point.

### CRISPRi Imaging Screen

The sgRNAs with the strongest phenotype from the pooled proliferation/survival screen were selected to generate individual sgRNA KD cell lines for *CEP290*, *CHD4*, *KMT2E*, *NSD1*, *OFD1*, *RFX3*, and *TAOK1*, as well as two non-targeting controls. CRISPRi cell lines were generated as previously described (H. R. Willsey et al. 2021; Sun et al. 2024), from hTERT dCas9 RPE1s and AICS dCas9 iPSC-derived NPCs. Knock-down was confirmed by qPCR using the ΔΔCT method (**Figure S3A**). RPE1 cells were plated on a 96-well glass bottom plate (Corning; Cat. no. CLS3603) at a density of 2×10^4^ cells per well. NPCs were plated on a matrigel-coated 96-well glass bottom plate at 4×10^4^ cells per well. RPE1s were serum starved (DMEM-F12 - FBS), then both RPE1s and NPCs were fixed after 24 hours in 4% paraformaldehyde. We permeabilized cells for 15 minutes in PBST (PBS, 0.2% Triton X-100) and blocked in blocking buffer (PBS, 0.2% Triton X-100, 2% BSA) for 45 minutes at room temperature. Cells were incubated in blocking buffer with primary antibody overnight at 4°C. ARL13B primary antibody (1:500, ProteinTech; Cat. no. 17711-1-AP) was used to visualize cilia. The cells were then washed three times in PBST for a total of 45 minutes and incubated for 1 hour at room temperature in a blocking buffer with goat anti-rabbit secondary antibody (1:1000, Fisher Scientific; Cat. no. A32732) as well as DRAQ5 (1:500, Fisher Scientific; Cat. no. 5016967). Stained cells were then washed three times in PBST for a total of 45 minutes, before being stored at 4°C in PBS for imaging. Images were acquired using a Zeiss 980 LSM confocal microscope with 20x and 63x objectives.

### CRISPRi Imaging Screen - Data Analysis

Cilia count was determined using the CellProfiler 4.2.5 software (McQuin et al. 2018). We adapted the ‘Speckle Counting’ pipeline to reliably identify cilia. First, the ARL13B channel was enhanced to remove background noise. BFP+ nuclei (positively transduced cells) were identified using a diameter range of 40-140 pixel units, threshold range of 0.0-1.0, threshold strategy set to ‘Global’, and threshold method set to ‘Minimum Cross-Entropy’. Cilia of these BFP+ cells were counted using a diameter range of 5-30 pixel units, a threshold range of 0.2-1.0, threshold strategy set to ‘Global’, and threshold method set to ‘Otsu’. Percent cilia measurements were normalized based on average BFP+ cell density of the non-targeting control sgRNA #1 (Figure 4) or unnormalized (Figure S4). Statistical significance was determined using Dunn’s multiple comparisons test in Graphpad (Prism).

Cilia length was determined using the CiliaQ plugin on Fiji (Hansen et al. 2021). Briefly, cilia length was quantified by inputting z-stacks of ARL13B and BFP channels into ‘CiliaQ Preparator’. Images were checked by eye for errors in cilia identification and co-localization with BFP+ cells. Errors in cilia identification were corrected using the ‘CiliaQ Editor’. Finally, cilia intensity and length were calculated using ‘CiliaQ V0.1.4’ with minimum cilium size (voxel) set to 20. Cilia length measurements were normalized based on average BFP+ cell density of the non-targeting control sgRNA and cilia intensity measurements were normalized based on average DRAQ5 fluorescence. Statistical significance was determined using Dunn’s multiple comparisons test in Graphpad (Prism). Note: We only quantified positively transduced, BFP+ cells for all CRISPRi assays.

### *Xenopus* Husbandry and Microinjections

Male and female wild-type *Xenopus laevis* and *Xenopus tropicalis* were maintained and cared for according to established IACUC protocols. Ovulation was induced in females using human chorionic gonadotropin (Sigma) according to (Sive, Grainger, and Harland 2000) before performing natural matings or *in vitro* fertilizations. Localization work was done in *X. laevis,* while knockdown was done in *X. tropicalis*.

Human *TAOK1* cDNA sequence (NM_020791.4) was cloned into the GFP vector (C-terminal tag) pcDNA3.1+ and injected at 20 pg per blastomere at the 4-cell stage, targeting the epidermis. Injected animals were imaged by confocal microscopy on a Zeiss 980 LSM with a 63x oil objective in confocal mode.

We generated a translation-blocking *taok1* morpholino (MO) (5′-TTGTTGACGGCATCCTGCTTCAG-3′) to disrupt *taok1* expression in *X. tropicalis* or a standard control morpholino (5’-CCTCTTACCTCAGTTA-CAATTTATA-3’) purchased from Gene Tools (Philomath, OR). For cilia phenotype analysis we injected 3.32 ng of *taok1* MO or standard control per embryo into one cell at the 4-cell stage using a Zeiss Stemi 508 microscope, Narishige micromanipulator, and a Parker Picospritzer III. Centrin-CFP RNA was injected at 50 pg per embryo. Animals were fixed and stained at stage 30. For heart/brain phenotyping, 8.3 ng of *taok1* MO or standard control along with a dextran tracer, was injected unilaterally at the two-cell stage for brain phenotyping, or in both cells at the two-cell stage for heart phenotyping.

### Xenopus tropicalis Cilia Phenotyping

Stage 30 *X. tropicalis* embryos were fixed using 4% PFA diluted in PBS. Immunostaining was performed according to (H. R. Willsey et al. 2018), with the omission of bleaching. Acetylated alpha-Tubulin primary antibody (1:1000, Sigma, Cat. no. T6793) along with goat anti-mouse Alexa Fluor 555 (1:500, LifeTech, Cat. no. A32727) conjugated secondary antibody were used to visualize cilia. Phalloidin (1:500, LifeTech, Cat. no. A22287) was added during secondary antibody incubation. Samples were mounted on glass slides (within an area enclosed by a ring of vacuum grease) with PBS and coverslipped. Images were acquired on a Zeiss AxioZoom V16 with a 1× objective or a Zeiss LSM980 confocal microscope with a 63× oil objective. Images were acquired as *z*-stacks at system-optimized intervals and processed in Fiji as maximum intensity projections. We imaged approximately 100 CFP+ MCCs across 3-4 tailbuds for both experimental and control conditions and classified cells as having no phenotype (>50 cilia), moderate phenotype (10-50 cilia), or severe phenotype (<10 cilia). We then compared the number of severe or moderate versus no phenotype MCCs in experimental and control conditions using a Chi-squared test in R.

### *Xenopus tropicalis* Heart/Brain Phenotyping

Heart and brain phenotyping was performed according to (Rosenthal et al. 2021). Stage 46 tadpoles were fixed with 4% PFA in PBS. Immunostaining was performed according to Willsey et al. 2018, with the omission of the bleaching step whenever phalloidin was included. Acetylated alpha-Tubulin primary antibody (1:500, Sigma, T6793) along with goat anti-mouse Alexa Fluor 488 (1:500, LifeTech; Cat. no. A32723) conjugated secondary antibodies were used to visualize the brain. Phalloidin (1:500, LifeTech, Cat. no. A22287) was used to visualize the heart (actin). Animals were imaged on a Zeiss AxioZoom V16 with 1X objective. Brain region size was calculated from stereoscope images of brain immunostainings using the freehand select and measure functions in Fiji. The injected side was compared to the uninjected side (internal control). These measurements were from two-dimensional images taken from a dorsal perspective and are a reflection of relative size differences, not a direct quantification of cell number. Heart ventricle size was measured using the freehand select and measure functions in Fiji. Quantitative differences in heart ventricle size were calculated by comparing mean surface area between control versus *taok1* MO injected embryos. For both brain and heart phenotyping, statistical significance was determined using unpaired Mann-Whitney rank sum tests in Graphpad (Prism).

## ACKNOWLEDGEMENTS

We thank Nolan Wong and UCSF LARC for animal care; Milagritos Alva and Juan Arbelaez for lab maintenance; Ashley Clement, Gigi Paras, Sonia Lopez, and Linda Chow for administrative support; Martin Kampmann and Avi Samelson for expert advice and sharing of reagents for the CRISPRi screen; Jeremy Reiter, Mia Konjikusic, and Yue Liufor assistance with RPE-1 cell culture and cilia analyses. The authors would like to thank all members of the Willsey Labs as well as Matthew State for their invaluable intellectual input and support.

## CONTRIBUTIONS

Conceptualization: NT, AJW

Formal Analysis: NT, MCL, SW, TJN

Funding Acquisition: NT, AJW, HRW

Investigation: NT, MCL, EK, EB

Methodology: NT, MCL, EK, EB, JD, AJW, HRW

Project Administration: TJN, AJW, HRW

Resources: EB, NS, TJN, AJW, HRW

Supervision: TJN, AJW, HRW

Validation: NT, SW, MCL, TJN, HRW

Visualization: NT, MCL, EK

Writing (original draft): NT

Writing (editing): NT, TJN, AJW, HRW

## DATA AVAILABILITY

ASD genetic data are available via Satterstrom *et al*. (2020). CHD genetic data are available via Jin *et al*. (2017). Shared risk ASD-CHD genes are available in Table S1 and were derived from Satterstrom *et al*. (2020), Jin *et al*. (2017), Rosenthal *et al*. (2021), and SFARI Gene (Banerjee-Basu and Packer 2010)

## Supplemental Figures

**Figure S1.**
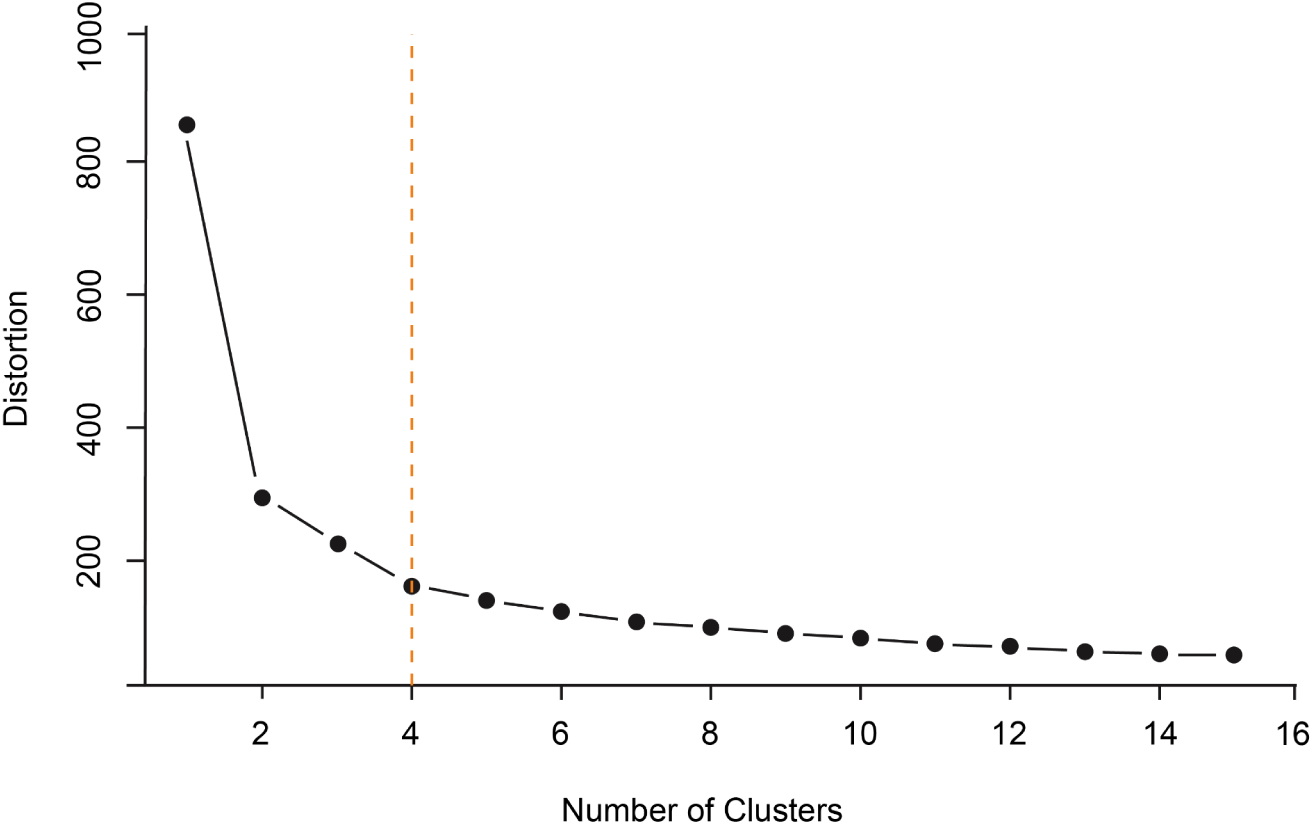
Elbow Plot. (A) Elbow method was used to determine the optimal clustering number for our 54 proliferation/differentiation screen genes with FDR < 0.1 and |Log2FC| ≥ 0.585 in at least 1 timepoint. We determined four to be the optimal cluster number, by identifying the inflection point of the graph (orange dotted line).

**Figure S2.**
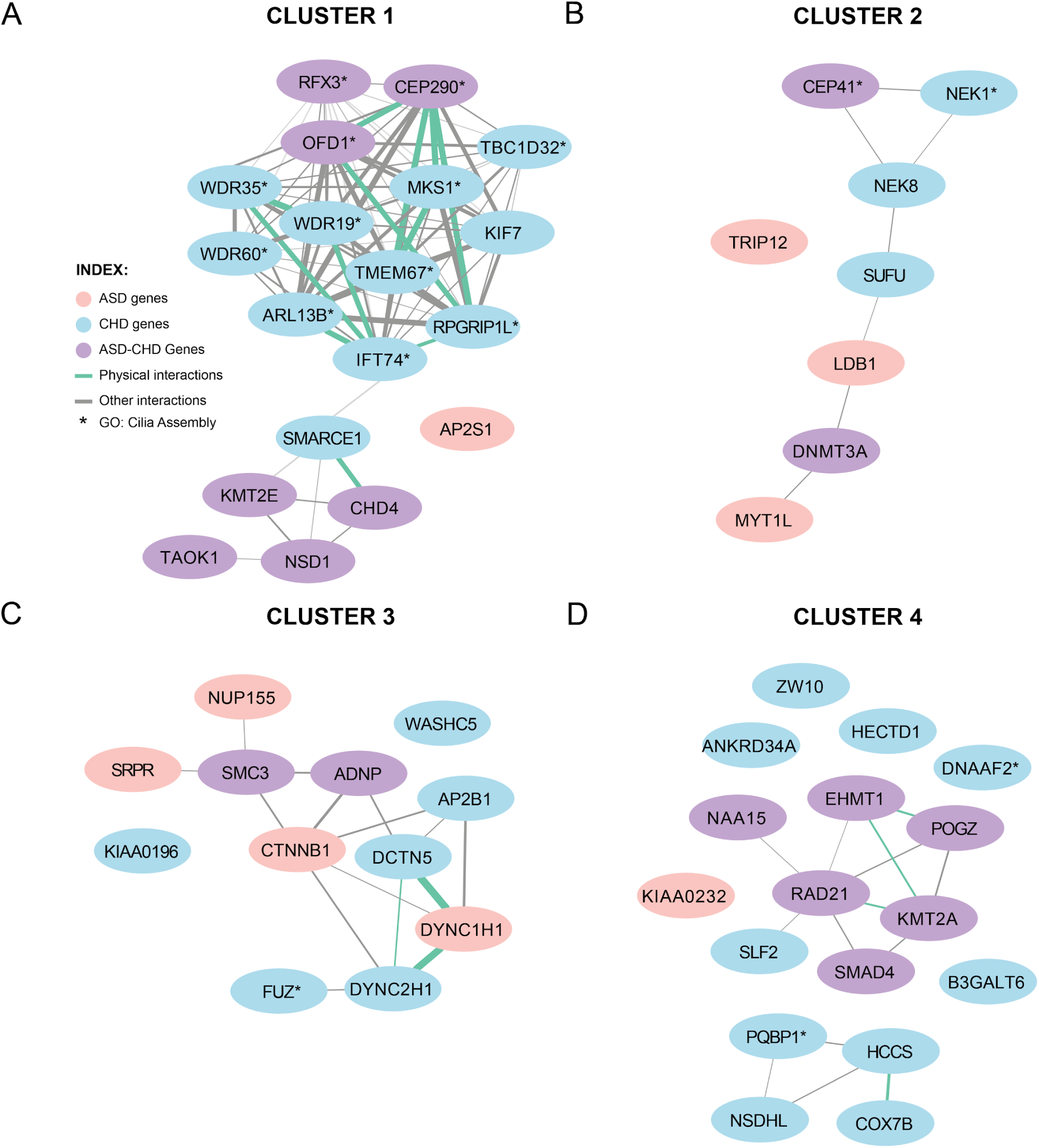
Cluster 1 is significantly enriched for interactions. (A-D) Visualization of Cluster 1-4 interactions identified from StringDB. ASD-genes (Satterstrom, 2019) are represented by a pink circle, CHD-genes (Jin, 2017) are represented by a blue circle, and predicted ASD-CHD genes are represented by a purple circle. Physical interactions are connected with a green line and all other types of interactions are represented by grey.

**Figure S3.**
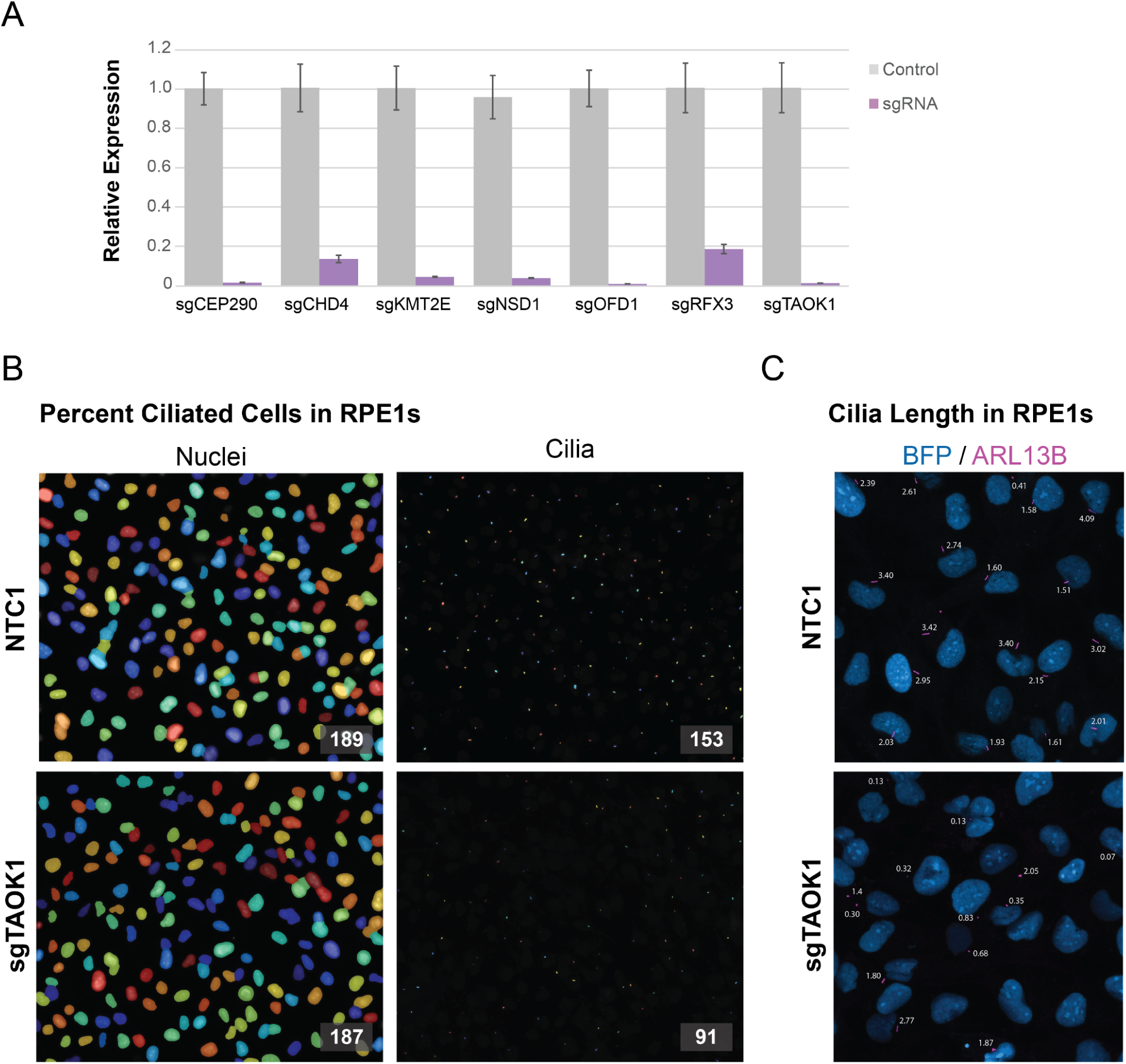
Cilia quantification in RPE1 cells. (A) Knockdown efficiencies of the 7 ASD-CHD gene sgRNAs were evaluated individually in established RPE1 cell lines by qPCR. (B) Representative image of percent cilia quantification of non-targeting sgRNA (NTC1; 81% ciliated) and sgTAOK1 (49% ciliated) using CellProfiler. (C) Representative image of cilia length quantification of non-targeting sgRNA (NTC1; Average length: 2.38µm) and sgTAOK1 (Average length: 0.98µm) using CiliaQ. *Image is represented as a 2D maximum projection, while cilia length was measured in 3D.

**Figure S4.**
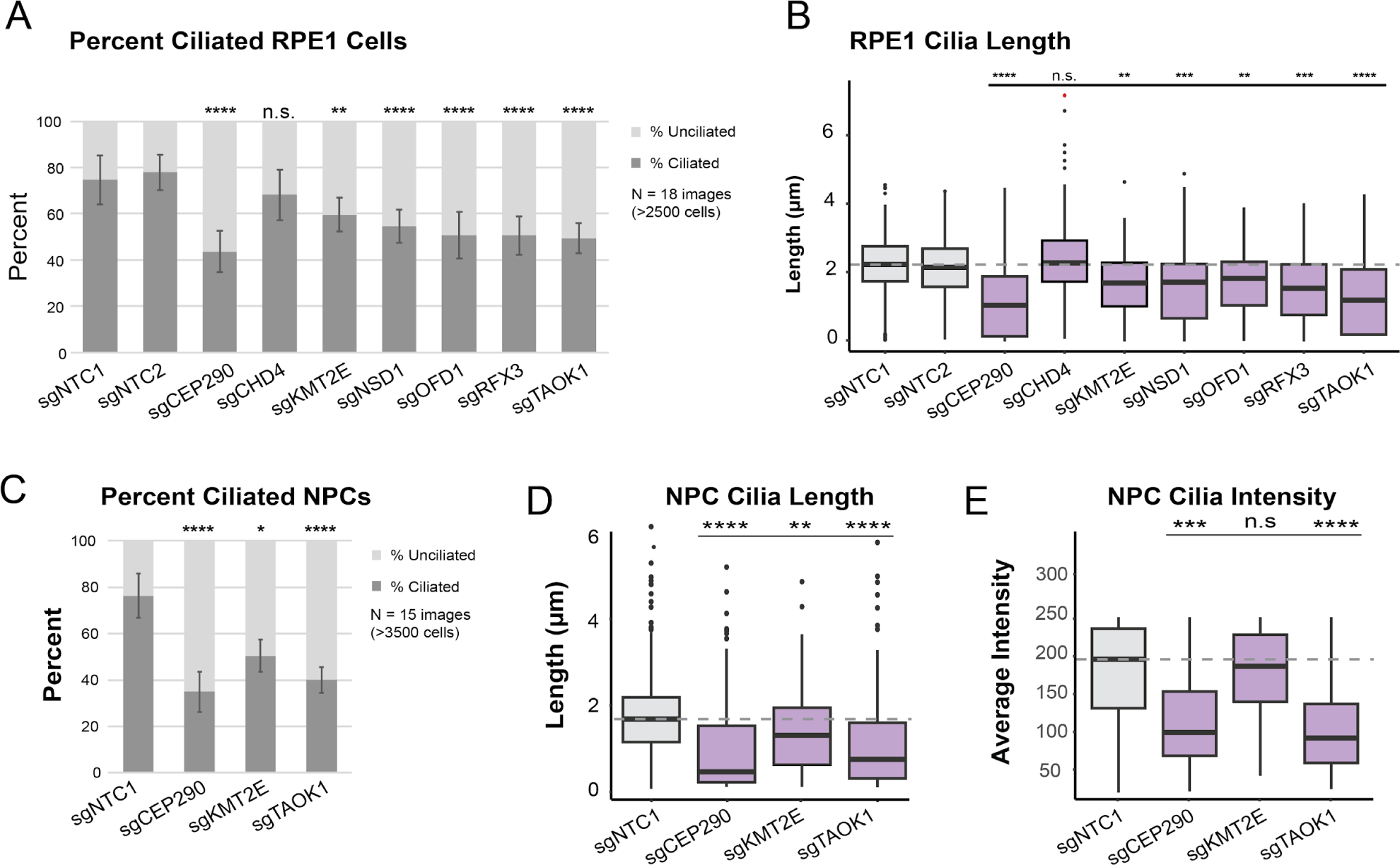
Knock-down of ASD-CHD genes disrupts primary cilia (without normalization) (A) We quantified percent ciliated cells in ≥2500 RPE1 cells across 18 images (3 biological replicates). (B) We captured ≥250 cells across 15 images (3 biological replicates) for quantification of cilia length (µm) in RPE1 cells. (C) We captured ≥3500 cells across 15 images (3 biological replicates) for quantification of percent ciliated cells in NPCs. (D) We captured ≥350 cells across 15 images (3 biological replicates) in NPCs. (E) Using the images from (F), we measured ARL13B intensity (a.u.) for quantification of cilia intensity in NPCs. \**Significance (Dunn’s multiple comparisons)*: *p < 0.05; \*\**p* < 0.01; ***p < 0.001; \*\*\*\**p* < 0.0001; n.s., not significant (*p* > 0.05)

**Figure S5.**
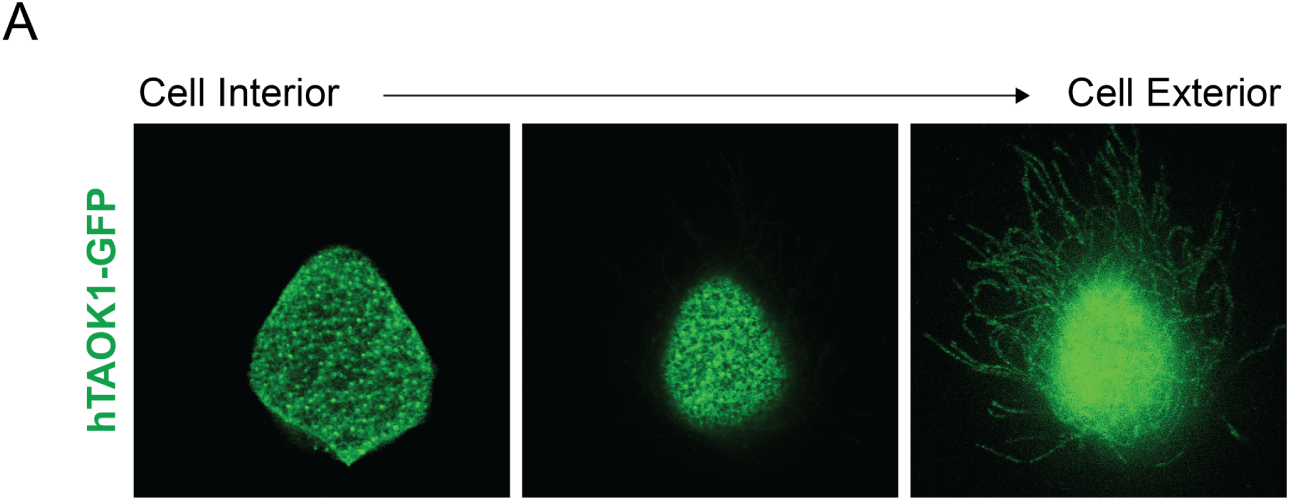
TAOK1 localizes to ciliary structures. (A) hTAOK1-GFP injected into *X. laevis* localizes to ciliary structures of epidermal multiciliated cells. Images depict a single cell across several focal planes from cell interior to exterior. hTAOK1-GFP appears to localize to basal bodies (left), actin (middle), and ciliary axonemes (right). These images are from an animal injected only with the GFP construct (so as not to have potential crosstalk from fluorescence channels from co-stains, but similar results were observed with costains for basal bodies and axonemes).

## Supplemental Tables

**<u>Table S1</u>** - List of screened genes ( Log2FC, PVal, ASD-satterstrom, CHD-Jin, ASD-CHD Rosenthal, ASD-CHD Genetic, CHD-SFARI)

**<u>Table S2</u>** - List of significant genes input for k-means clustering (per replicate, Cluster number, Log2FC, PVal, Category (ASD, CHD, ASD-CHD))

*Legend: Dx = day number; Rx = replicate number

**<u>Table S3</u>** - ToppGene Enrichments (Biological Process and Cellular Component) of Cluster 1 genes with CRISPRi screen genes used as background correction (Tab 1 - CRISPRi BG), as well as, default background correction (Tab 2 - Default BG)

